# Influence of non-metabolic microbial growth promotors (cAMP – activators) on the sensitivity to antimicrobials in the actually multiresistant microbial strains

**DOI:** 10.1101/143438

**Authors:** Artur Martynov, Tatyana Osolodchenko, Boris Farber, Sophya Farber

**Affiliations:** 14-Puskhinskays St., Kharkov 61057, Ukraine; 14-Puskhinskaya St., Kharkov 61057, Ukraine; 1781 East 17th St, #d6, Brooklyn, New York 11229, USA; 1323 E 18th St, Brooklyn, NY 11230, USA

**Keywords:** cAMP-inducers, antimicrobials, MDR strains, *Pseudomonas aeruginosa*, *Acinetobacter baumanii*, *Klebsiella pneumonia*, bacterial growth, sensitivity to antimicrobials

## Abstract

**Introduction:** The control over multi-resistant nosocomial strains of microorganisms has been becoming increasingly urgent in recent years. We suggest a new paradigm that eliminates killing or inhibiting the growth of bacteria. Excluding bacteria death supresses the selection of resistant strains of microorganisms. We have developed such non-metabolite growth promoters, which in very low doses stimulate the rapid growth of many bacteria strains. The mechanism of action of the enhancers is caused by the activation of the cAMP high doses accumulation process in the microbial cells. cAMP itself is a substrate for phosphorylation including DNA polymerases.

**Materials and methods:** The susceptible culture collection resistance strains *Pseudomonas aeruginosa MDR Kharkov IMI1, Acinetobacter baumanii MDR Kharkov-IMI1*, and *Clebsiella pneumonia MDR Kharkov-IMI1* were used. The following antimicrobial agents of known potency were evaluated: ciprofloxacin, polymyxin, amikacin. The same broth, but containing 0.001% enhancers (under patenting), has been used for further passaging for MDR strains. Characteristics of bacterial growth were determined in a medium compared at the control group – the broth without enhancers.

**Results and discussion:** Enhancers contribute to a significant increase in the antimicrobial sensitivity to polymyxin, ciprofloxacin and amikacin in multi-resistant strains of bacteria. Changes in the growth characteristics and antimicrobial sensitivity are observed only in the second passage that demonstrates the need for the further studies of the molecular mechanisms of the cAMP effect on the division and growth of microbial cells.

## Introduction

The fight against multiresistant nosocomial microorganisms is becoming increasingly important in recent years [1].

This is due to the uncontrolled use of antibiotics at home, as well as the massive use of antiseptics in hospitals. Recent studies of bacterial resistance mechanisms call attention to enhancement of protective factors like biofilm formation and non-growing state by bacteria [2].

Currently available antibiotics are intended mainly to fight against vegetative forms of microorganisms. This is responsible for low efficacy of antibiotics in treatment of pathologies caused even by highly susceptible microbial strains, but with a high degree of biofilm formation [3].

In addition, the uninterrupted selection of multiresistant strains in hospitals and at home setting under the influence of antibacterial drugs, the mandatory use of antibiotics in veterinary medicine in animal rearing for meat and milk production also activates the selection of multiresistant strains, even in circumstances when one does not get sick, but only consumes milk or meat, saturated with sub-lethal doses of antibiotics [4,5].

Based on the above, the classical approaches associated with the development of new antibiotic lines are gradually losing their relevance. It is necessary to change the paradigm of combating multi-drug resistance of microorganisms [6]. We offer a new paradigm that eliminates killing of bacteria or inhibition of their growth. Excluding bacterial death excludes the selection of resistant strains of microorganisms.

A fact known from industrial microbiology: toxins production by bacteria in industrial conditions is observed only in the starving nutrient media containing aggressive factors (serum, red blood cells, the heart or the brain extracts), but not rich in carbohydrates. This fact is known for bacteria such as C. diphtheria, P. aeruginosa, tetanus and pertussis bacillus. [7]. If these bacteria are transferred to the rich, non-starvation medium without aggressive factors, they begin growing rapidly, but without synthesizing toxins in general [8,9], besides, they stop synthesizing biofilm and other resistance factors [10].

The phenomenon following from the facts above: if the bacteria do not fight for their existence with external aggressive factors (in our case - host immune system), they become harmless to the organism. To convert them to a state of “euphoria”, they need to be cheated - to be shown that the aggressive factors of the environment are absent, and that there is an urgent need to start active division. We have developed such non-metabolite growth promoters, which stimulate the rapid growth of almost all bacterial strains in very low doses [11,12].

The mechanism of action of the enhancers is caused by activation of accumulation process of high cAMP doses in microbial cells. cAMP itself is a substrate for phosphorylation, including DNA polymerases. Their activity is increased several hundred times after phosphorylation. The rapid bacterial growth is completely incompatible with the release of majority of acquired virulence factors (including lactamases), toxin production, film forming [13].

At first, bacteria “clear the area” by excreting toxins, and start dividing rapidly only when the enemy is killed. So we got an idea of using antibacterials after growth stimulation – in this case, the bacteria is supposed to lose its acquired resistance in the process of rapid growth and even become harmless to the body and the immune system.

We have previously studied more than 200 cAMP activators and their various combinations with each other in order to determine the most active. Only one of the combinations studied showed a significant acceleration of bacterial growth.

The goal of the study was investigation of the effect of cAMP accumulation activators on the antibiotic resistance for the most multiresistant *P. aeruginosa* [14], *A. baumanii* [15, 16] and *K. pneumonia* [17] strains. These strains were isolated in Kharkiv hospitals and delivered to our institute for research. They were resistant to all antibiotics and antimicrobials. Only *Pseudomonas aeruginosa* remained slightly sensitive to polymyxin in 10-fold MIC.

## Materials and methods

### Strains and antibiotics

The susceptible culture collection strains *Pseudomonas aeruginosa MDR Kharkov IMI1, Acinetobacter baumanii MDR Kharkov-IMI1*, and *Clebsiella pneumonia MDR Kharkov-IMI1* were used. The following antimicrobial agents of known potency were evaluated: ciprofloxacin (Bayer AG, Wuppertal,Germany), polymyxin (Xellia Pharmaceuticals ApS, Denmark), amikacin (Arterium, Kiev, Ukraine). Antibiotics were dissolved in water at MICs in Mueller-Hinton broth (MHB) (table 1 for ATCC strains). All subsequent dilutions were made in cation-supplemented Mueller-Hinton broth (Difco Laboratories, USA) and prepared fresh for each experiment. The same broth, but containing 0.001% enhancers: (under patenting), was used for further passaging for MDR strains. Bacterial growth characteristics were determined in a medium compared against the control group - broth without enhancers. Enhancers are FDA-approved water soluble substances that have been used in medicine as drug products for a long time, but for other purposes. Taken separately from each other, they are unable to stimulate bacterial growth and have no influence on multidrug resistance.

In our studies, we used antibacterials at MICs for strains presented as line in the table 1-9. All test MDR strains were resistant to all studied antimicrobials at standard MICs [18]. The inoculum was added at initial concentration of 5 x 10^3^ CFU/ml from an exponential-phase culture. Each passage included the tubes incubated for 72 h at 37°C, after which the samples (10 μl) were transferred from tubes onto blood agar plates which were incubated for 18 h at 37°C for CFU counting by live cells. Also, cell numbers were determined from an optical density-CFU standard curve by Densi-LA-meter (ERBA Lachema, Czech Republic) after incubation for 72 h in each passage. In parallel, samples (10 μl) were transferred to the new tubes as next passage. Passage-killing curves were plotted using the techniques described above. The average value of absorbance in liquid and solid medium was used for each CFU/ml point (including live and killed microbial cells).

Results were processed using the analysis of variance.

**Fig. 1.**
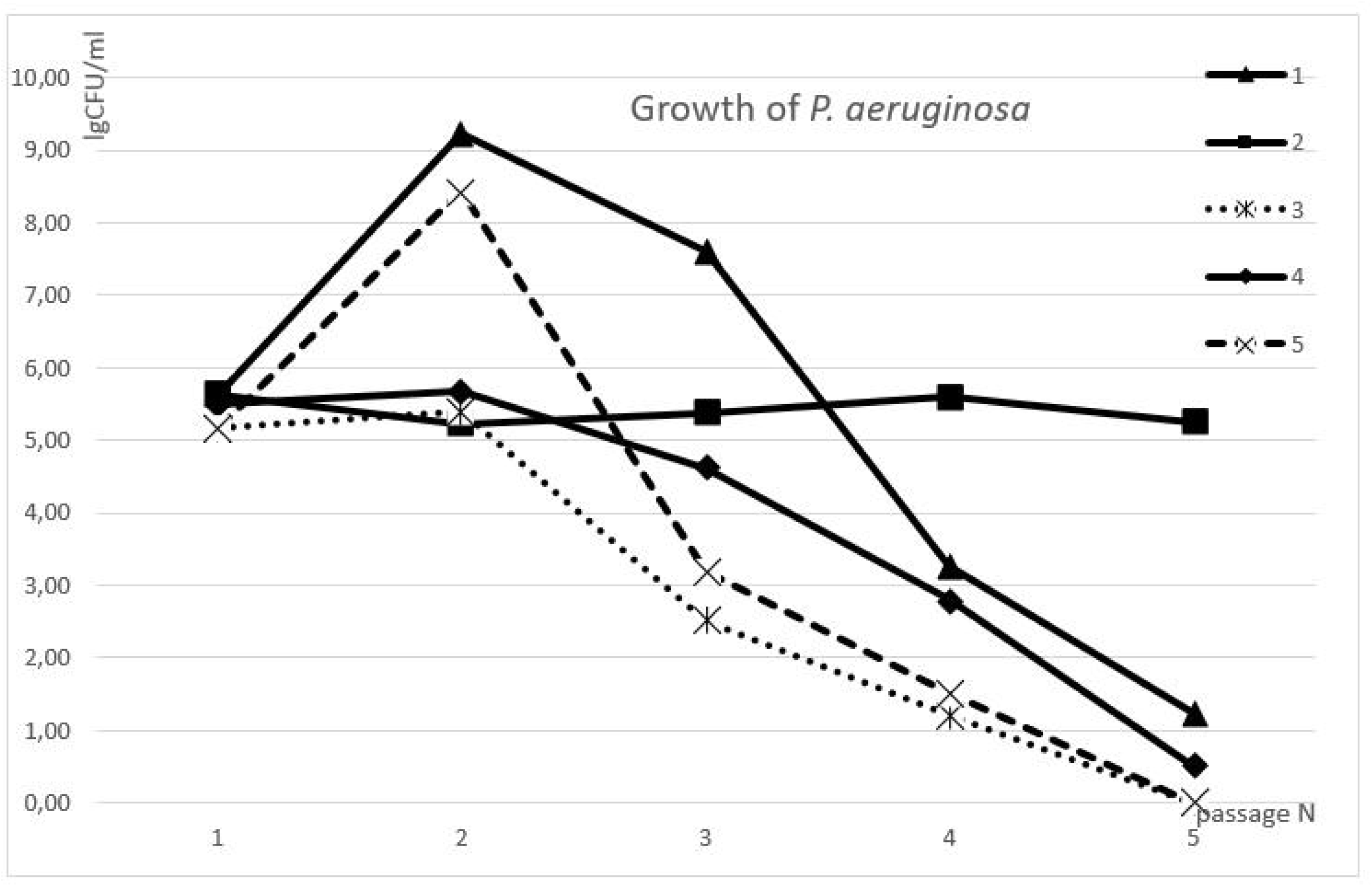
Relation between the passage number of MDR strain *P. aeruginosa* and lg CFU/ml parameter in different media and with different antibacterials: 1 –growth on MHB with enhancers; 2 – growth on MHB without enhancers; 3 – growth on MHB with enhancers and polymyxin (0.5 μg/ml); 4 – growth on MHB with enhancers and ciprofloxacin (1.0 μg/ml); 5 – growth on MHB with enhancers and amikacin (8.0 μg/ml).

**Table 1.** Antibacterial activities of some antibacterials in the ATCC and MDR-strains

| N | Microorganism | Antimicrobials | MIC (µg/ml) | MBC (µg/ml) |
| --- | --- | --- | --- | --- |
| 1 | *Pseudomonas aeruginosa*<br>ATCC 27853 | polymixine | 0,5 | 3,0 |
| 2 |  | ciprofloxacin | 1,0 | 24,0 |
| 3 |  | amikacin | 8,0 | 24,0 |
| 4 | *Acinetobacter baumannii* ATCC<br>19606 | polymixine | 0,5 | 1,0 |
| 5 |  | ciprofloxacin | 1,0 | 24,0 |
| 6 |  | amikacin | 8,0 |  |
| 7 | *Clebsiella pneumoniae* ATCC<br>13883 | polymixine | 0,5 | 3,0 |
| 8 |  | ciprofloxacin | 1,0 | 3,0 |
| 9 |  | amikacin | 8,0 | 24,0 |
| 10 | *Pseudomonas aeruginosa* MDR<br>Kharkov IMI1 | polymixine | 32,0 | 128,0 |
| 11 |  | ciprofloxacin | - | - |
| 12 |  | amikacin | - | - |
| 13 | *Acinetobacter baumannii* MDR<br>Kharkov IMI1 | polymixine | - | - |
| 14 |  | ciprofloxacin | - | - |
| 15 |  | amikacin | - | - |
| 16 | *Clebsiella pneumoniae* MDR<br>Kharkov IMI1 | polymixine | 64,00 | - |
| 17 |  | ciprofloxacin | - | - |
| 18 |  | amikacin | - | - |

### Results and discussion

As can be seen in Fig. 1, the growth curves of *P. aeruginosa MDR* in 5 passages on the medium without enhancers are almost identical (line 2): cell number increase is observed after inoculation of 10^3^CFU/ml to 10^5^CFU/ml at each passage. Initially low concentration of inoculum (10^3^CFU/ml) was taken in view of preliminary data on the potential bacterial growth within 72 hours.

**Fig. 2.**
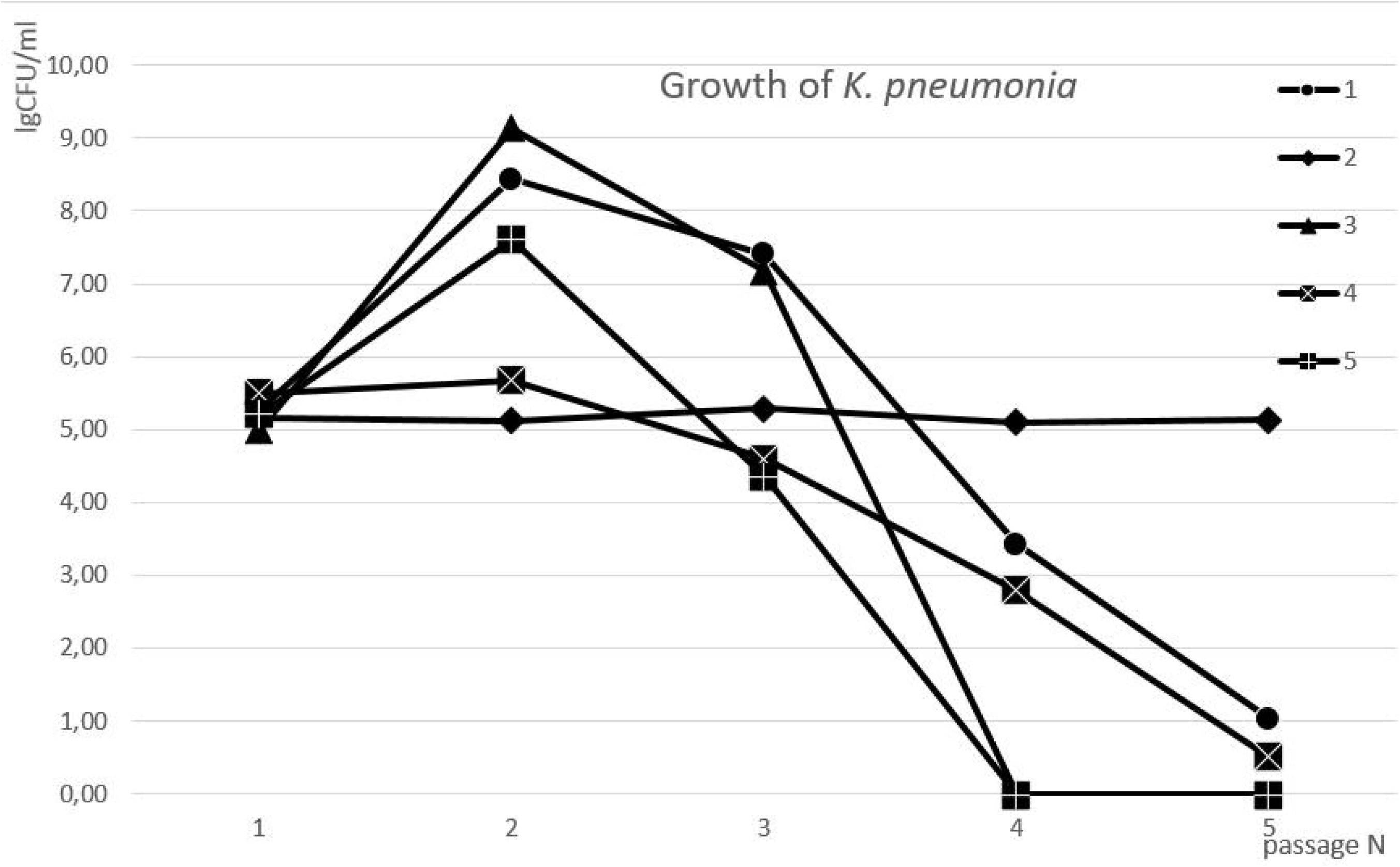
Relation between passage number of MDR strain *K. pneumonia* and lg CFU/ml parameter in different media and with different antibacterials: 1 – growth on MHB with enhancers; 2 – growth on MHB without enhancers; 3 – growth on MHB with enhancers and polymyxin (0.5 μg/ml); 4 – growth on MHB with enhancers and ciprofloxacin (1.0 μg/ml); 5 – growth on MHB with enhancers and amikacin (8.0 μg/ml).

*P. aeruginosa* growth in media supplemented with enhancers is statistically different from the growth medium without enhancers: as shown in the figure (line 1), at the second passage, the CFU number was already four orders of magnitude higher versus the control; at the third passage, growth rate was markedly reduced to 10^7^ CFU/ml. At passages 4-5, inhibition of bacterial growth in the presence of enhancers was observed, which required further molecular biological studies of this phenomenon. It is also not quite clear why the growth at the first passage showed almost no differences from the control 5.62 ± 0.03 lg CFU/ml with the rapid growth enhancement at the second passage.

Polymyxin addition to the medium with enhancers at concentration 0.5 μg/ml also resulted in significant changes in the microbial growth dynamics. At the first passage, the differences versus the control group were actually absent. Also, virtually no increment was observed at the second passage (line 3) (5.40 ± 0.35 lg CFU/ml). Previously multidrug resistant *P. aeruginosa* strain became sensitive to polymyxin at the passage 3-4. At passage 5, no bacterial growth at all was observed in the presence of polymyxin and no live bacteria were identified after blood agar inoculation. At passages 3-5, the differences from control (line 2) were statistically significant (P <0.05), as well as differences between the lines 1 and 3.

A similar growth curve was observed with the addition of 1 μg/ml ciprofloxacin to the medium. Initially, the bacterium was resistant to both polymyxin and ciprofloxacin and did not respond to the presence of both antibacterial drugs in the medium in the recommended doses of 0.5 and 1.0 μg/ml, respectively. Growth and sensitivity to ciprofloxacin at the first passage were not statistically different from the control, whereas ciprofloxacin significantly inhibited bacterial growth already at the second passage (no statistically significant increment was observed versus the medium with enhancers). At the third passage, statistically significant (P <0.05) inhibition of bacterial growth (4.6 ± 0.13 lg CFU/ml) was observed versus both the control without enhancers (line 2), and versus the control group - the medium with enhancers (line 3). At passage 4-5, the bacterium was already highly sensitive to ciprofloxacin. Subsequent inoculation of the bacteria onto blood agar in the presence of disks with antibiotics has confirmed the loss of MDR by the bacteria and its high sensitivity to polymyxin and ciprofloxacin.

Amikacin addition in the dose 8 μg/ml to the culture medium with enhancers also led to a change in the survival curve upon passaging. The same as in the first two cases with polymyxin and ciprofloxacin, the first passage with amikacin was not different from the control – the increment of 2 orders of magnitude was the same as for the controls. The second and third passages were not statistically different from the line 1 with enhancers, whereas no bacterial growth was observed at all at passages 4 and 5, and the bacteria could not be revived on blood agar. In fact, dramatic biochemical changes were observed at passages 3-4.

As can be seen in Fig. 2, *K. pneumonia* growth curves at 5 passages in the medium without enhancers were almost identical (line 2): increase in the number of cells by 2 orders of magnitude with inoculum from 10^3^ CFU/ml to 10^5^ CFU/ml was observed at each passage. Growth of *K. pneumonia* in the medium with enhancers was statistically significantly different from the growth in the medium without enhancers (8.43 ± 0.40 lg CFU/ml). As can be seen from the figure (line 1), CFU number at the second passage was already 4 orders of magnitude higher than in the control; at the third passage, the growth rate was markedly reduced to 10^7^ CFU/ml. At passages 4-5, inhibition of bacterial growth was observed in the presence of enhancers; this fact requires further molecular-biological studies of this phenomenon. Addition of 0.5 μg/ml polymyxin to the culture medium with enhancers also resulted in significant changes in the growth dynamics: while the differences were absent in the first passage, the increment at the second passage (line 3) was even higher than in the control medium with enhancers (10^9^ CFU/ml). MDR *K. pneumonia* growth at passage 3 hardly differed from the antibiotic-free control; no bacterial growth was observed in the presence of polymyxin at passages 4-5, and no live bacteria were identified after inoculation into blood agar. At passages 4-5, the differences from the control (line 2) were statistically significant (P <0.05).

After 1 μg/ml ciprofloxacin addition in the medium, the pattern quite different from *Pseudomonas aeruginosa* was observed. Initially, the bacterium was resistant to both polymyxin and ciprofloxacin, and did not respond to the presence of both antibacterial drugs in the recommended doses of 0.5 and 1.0 μg/ml, respectively. Microbial count increment and sensitivity to ciprofloxacin at the first passage were not statistically different from the control, whereas ciprofloxacin already significantly inhibited bacterial growth at the second and third passages (P <0.05): 5.67 ± 0.37 lg CFU/ml and 4.60 ± 0.13 lg CFU/ml, respectively. The fourth passage showed statistically significant (P <0.05) bacterial growth inhibition (2.80 ± 0.30 lg CFU/ml) versus both the control without enhancers (line 2) and the control group – medium with enhancers (line 3). The bacterium was already highly sensitive to ciprofloxacin, and yielded single colonies on blood agar at the passage 5. Subsequent inoculation into blood agar containing disks with antibiotics has confirmed the bacteria’s MDR loss and high sensitivity to polymyxin and ciprofloxacin.

Addition of 8 μg/ml amikacin to the medium with enhancers also led to a change in the survival curve upon passaging. The first and second passages did not differ from controls. The third passage was statistically different from the line 1 with the enhancers (experimental data: 4.37 ± 0.47 lg CFU/ml), whereas in the passage 4 and 5, no bacterial growth was observed at all, and bacterial revival on blood agar was not successful. In this case, we can see the regularity typical of multiresistant Pseudomonas aeruginosa as well. Probably, such an abrupt transition to the bactericidal effect is related with amikacin mechanism of action.

**Fig. 3.**
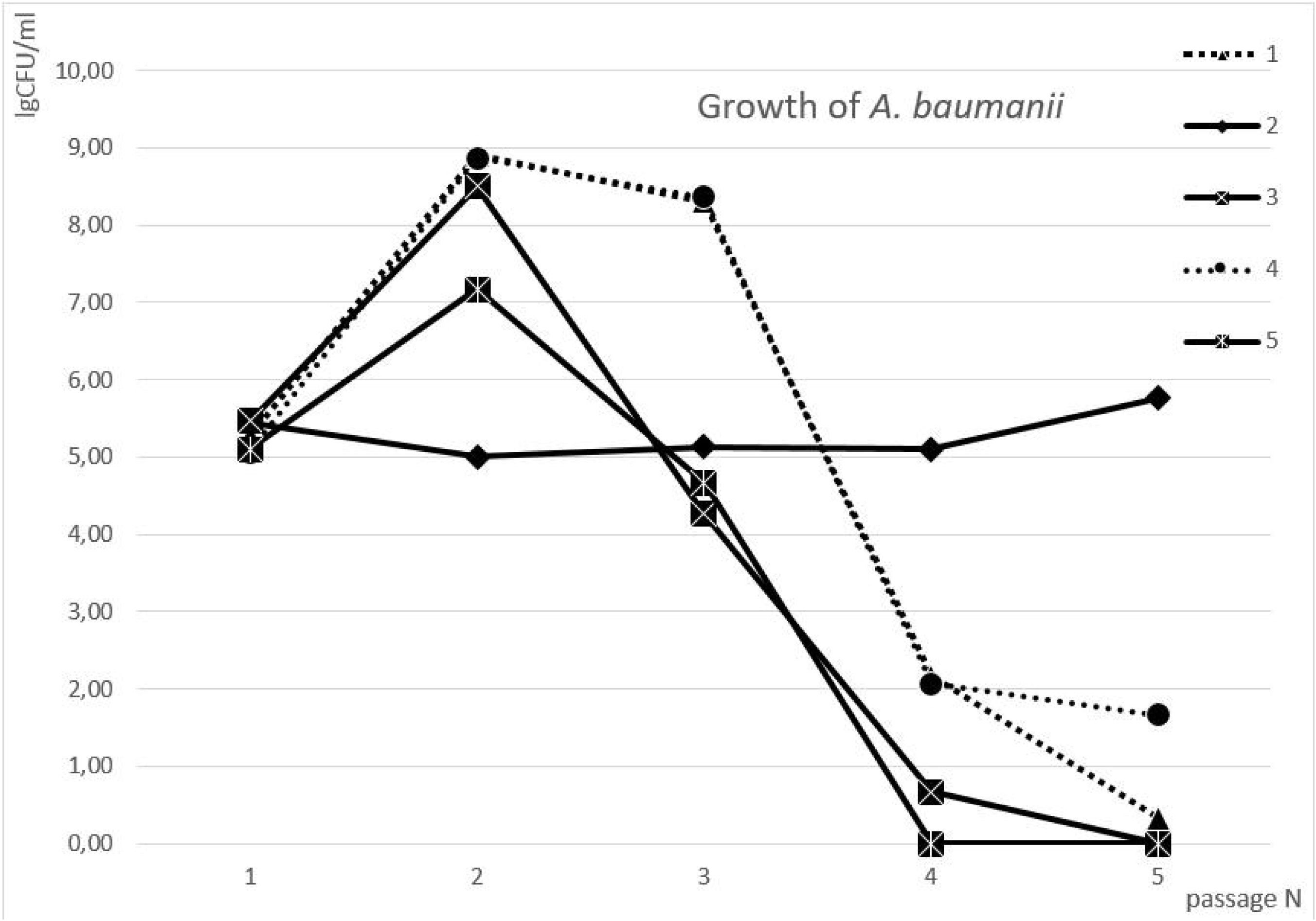
Relation between passage number for *A. baumanii* MDR strain and lg CFU/ml parameter in different media and with different antimicrobials: 1 – growth on MHB with enhancers; 2 – growth on MHB without enhancers; 3 – growth on MHB with enhancers and polymyxin (0.5 μg/ml); 4 – growth on MHB with enhancers and ciprofloxacin (1.0 μg/ml); 5 – growth on MHB with enhancers and amikacin (8.0 μg/ml).

As can be seen from Figure 3, *A. baumanii* growth curves at 5 passages on the medium without enhancers are almost identical (line 2): increase in the number of cells by 2 orders of magnitude with inoculum from 10^3^ CFU/ml to 10^5^ CFU/ml was observed at each passage (Fig.3). *A. baumanii* growth on the medium with enhancers was statistically significantly different from the growth on the medium without enhancers (8.90 ± 0.10 lg CFU/ml). As can be seen from the figure (line 1), CFU number at the second passage was already 4 orders of magnitude higher than in the control, but at the third passage, the growth rate was significantly reduced to 10^8^ CFU/ml. Bacterial growth inhibition in the presence of enhancers was observed at passages 4-5, which requires further molecular biological studies of this phenomenon. Addition of 0.5 μg/ml polymyxin to the medium with enhancers also resulted in significant changes in growth dynamics, beginning from passage 3. The growth of *A. baumanii* MDR strain at the passage 3 was statistically significantly lower than the control (4.27 ± 0.25 lg CFU/ml); bacterial growth at passage 4 in the presence of polymyxin was decelerated (0.67 lg CFU/ml), and no growth at all was observed at passage 5. No live bacteria could be identified at passage 5 on blood agar. Differences from the control (line 2) were statistically significant (P <0.05) at passages 3-5.

Addition of 1 μg/ml ciprofloxacin to the medium showed actually no statistically significant difference from the line 1, indicating *A. baumanii* lack of susceptibility to ciprofloxacin.

Addition of 8 μg/ml amikacin to the medium with enhancers led to a change in the survival curve along all passages. The first and second passages were slightly different from the control. The third passage was statistically different from the line 1 with the enhancers (experimental data: 4.67 ± 0.15 lg 4 CFU/ml), whereas no bacterial growth at all was observed at passage 4 and 5, and bacterial revival on blood agar was not successful. In this case, we observe the regularity typical of multiresistant *Pseudomonas aeruginosa* and *Klebsiella pneumonia*. Probably, such an abrupt transition to the bactericidal effect is related with amikacin mechanism of action.

## Conclusion

1. It has been established as a result of investigation of the influence of cAMP accumulation activators (enhancers) on multidrug-resistant microorganisms (*P. aeruginosa, K. pneumonia, A. baumanii*) that rapid growth stimulation by enhancers at the second passage is ended up in all cases with statistically significant inhibition of bacterial growth at subsequent passages until complete cessation of growth at passages 4-5.
2. Enhancers contribute to a significant increase in sensitivity of multiresistant bacterial strains to antimicrobials (polymyxin, ciprofloxacin and amikacin).
3. Changes in growth characteristics and antimicrobial sensitivity are observed only since the second passage, which demonstrates the need for further study of the molecular mechanisms of cAMP effect on the microbial cells division and growth.

## Declarations

Conflict of interest

Authors declare that they have no conflict of interests to disclose.

## Acknowledgements

This work was supported in part by the Farber Center of Academic Success (New-York, USA), “Noigel LLC.” (New-York, USA), the National Academy of Medical Sciences of Ukraine.

